# Influenza incidence prediction for the United States: an update for the 2018-2019 season

**DOI:** 10.1101/411348

**Authors:** Xiangjun Du, Yousong Peng, Mi Liu, Mercedes Pascual

## Abstract

**Introduction:** Seasonal influenza causes a high disease burden every year in the United States and worldwide. Anticipating epidemic size ahead of season can contribute to preparedness and more targetted control and prevention of seasonal influenza.

**Methods:** A recently developed process-based epidemiological model that incorporates evolutionary change of the virus and generates incidence forecasts for the H3N2 subtype ahead of the season, was previously validated by several statistical criteria, including an accurate real-time prediction for the 2016-2017 influenza season. With this model, a new forecast is generated here for the upcoming 2018-2019 season. The accuracy of predictions published for the 2017-2018 season is also retrospectively evaluated.

**Results:** For 2017-2018, the model correctly predicted the dominance of the H3N2 subtype and its higher than average incidence. Based on surveillance and sequence data up to June 2018, the new forecast for the upcoming 2018-2019 season indicates low levels for H3N2, and suggests an H1N1 dominant season with low incidence of influenza B.

**Discussion:** Real-time forecasts, those generated with a model that was parameterized based on data preceding the predicted season, allows valuable evaluation of the approach. Anticipating the dominant subtype and the size of the upcoming epidemic ahead of season informs disease control. Further studies are needed to promote more accurate ahead-of-season forecasts and extend the approach to multiple subtypes.

---

Human infections caused by the seasonal influenza virus impose a large burden on public health in the United States and worldwide. Anticipating the severity of the upcoming influenza season can contribute to timely and effective preparation, including targeted resource allocation and vaccination campaigns. To predict influenza incidence for the H3N2 subtype in the United States, a process-based model (EvoEpiFlu) was recently developed that incorporates evolutionary information into epidemiology based on the mathematical SIRS formulation of transmission dynamics (for Susceptible, Infected, Recovered and Susceptible immune classes in the population) (*1,2*). The SIRS model (without evolutionary change) is being used successfully to implement within-season forecasting (*3*). EvoEpiFlu makes additional use of evolutionary information related to antigenic change and based solely on readily available genetic sequences of the virus, to extend the lead time of prediction and make it possible to produce forecasts ahead of the influenza season (*1,2*). There are two versions of the EvoEpiFlu model based respectively on either a continuous or a discrete covariate for evolutionary change. The ‘continuous’ version of the model relies on an evolutionary index measuring how much the genetic sequences encoding antigenic sites have changed relatively to the recent past. The ‘discrete’ or ‘cluster’ version relies instead on the proportion of antigenic variants (PAV) inferred on the basis of an existing genotype-phenotype map (*4*), to identify intermittent periods of antigenic change (*1,2*). An increasing and high value of either measure of evolutionary change indicates the emergence of novel variants and a potential for high levels of future infection. A decreasing trend for PAV implies similar circulating strains and potentially fewer infections due to immune protection (*1,2*). Similarly, a decrease from sustained high values of the evolutionary index would preclude a high number of susceptible individuals and reduce transmission (*1,2*).

Based on the EvoEpiFlu model, we had previously generated a prediction for the 2017-2018 influenza season before it arrived, which we can now be validated with the updated surveillance data (Figure 1, bottom panel). The prediction was a high H3N2 season with an estimated higher than average incidence rate (proportion of the population infected) of 0.09 (median, with 95% confidence interval [0.05,0.14]) (Figure 1, dotted black curve and shaded black area, bottom panel). For comparison, the observed average incidence rate of H3N2 for the past sixteen years is 0.06. Our prediction is consistent with an observed value of 0.14 for the season that just ended (*5*), with the actual rate higher than median and within, but at the higher limit, of our prediction interval (Figure 1, bottom panel, solid black curve).

**Figure 1.**
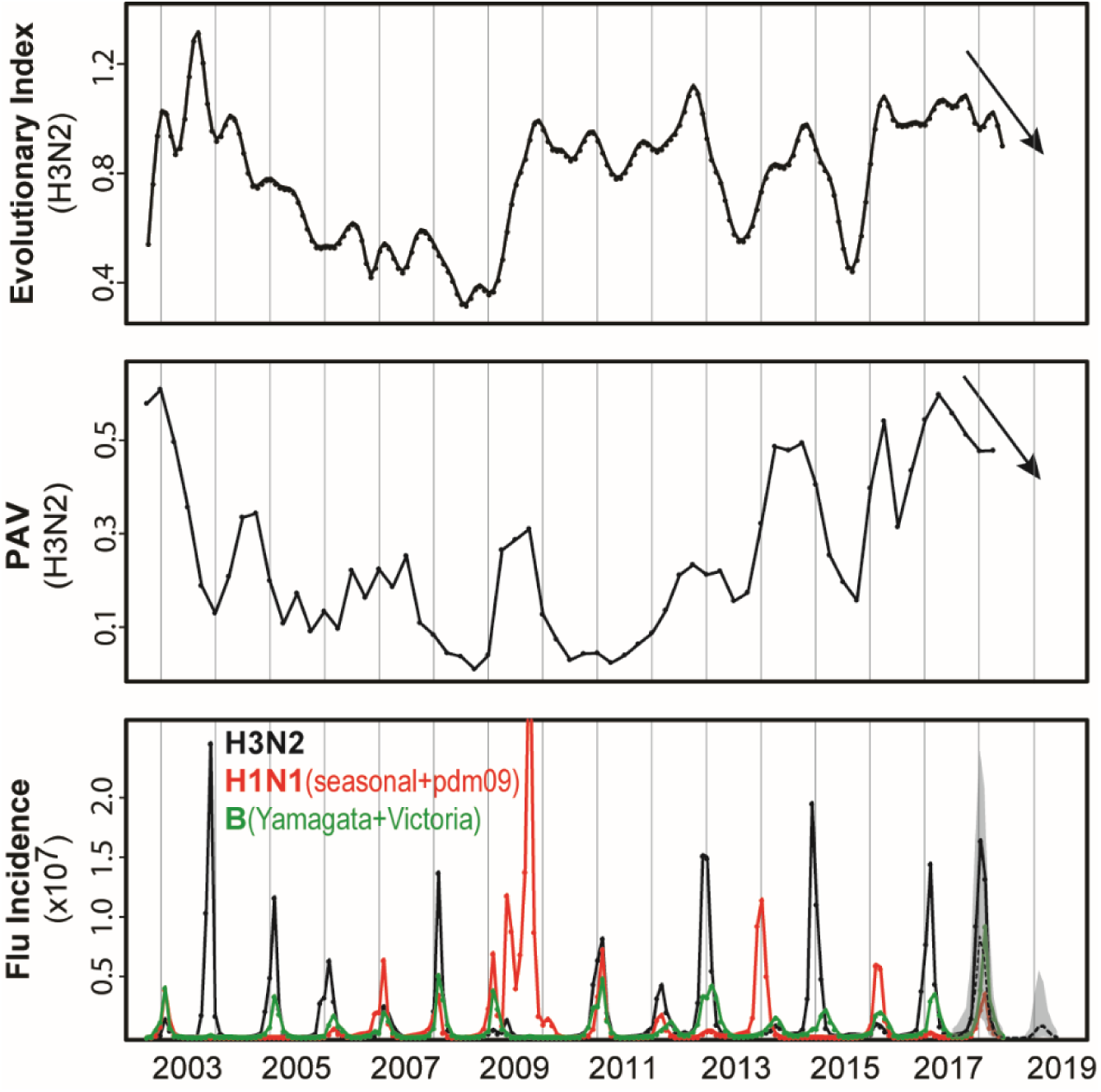
Evolutionary components and forecasts of the EvoEpiFlu model. Top panel: the evolutionary index calculated monthly. Middle panel: proportion of antigenic variants (PAV) calculated for strains in each quarter compared to strains in the past 12 months. Bottom panel: incidence and forecasts. The monthly observed incidence data of seasonal influenza in the United States are shown, with H3N2 in black, H1N1 (both seasonal and pandemic H1N1) in read and B (both Yamagata and Victoria lineages) in green. H3N2 incidence forecasts for 2017/2018 and 2018/2019 seasons are shown in dotted lines (with shaded 95% uncertainty intervals) based on data up to June, 2017 and 2018 respectively.

Encouraged by the prediction skill of EvoEpiFlu in this last season and before (*1,2*), we report here a routine forecast for the upcoming 2018-2019 influenza season. Both the PAV and the Evolutionary Index, exhibit a decreasing trend (Figure 1, top 2 panels). On the basis of this trend in PAV and the incidence data up to June 2018, EvoEpiFlu’s cluster model predicts that the upcoming 2018-2019 season will be H3N2 low, with an estimated median incidence rate of 0.013 ([0.003,0.041], 95% confidence interval) (Figure 1, bottom panel, black dotted curve and shaded area). The continuous model gives similar result with an estimated median incidence of 0.059 ([0.022,0.109], 95% confidence interval), at the edge of the historical average.

Because EvoEpiFlu is specifically designed for H3N2, it is not yet possible to apply it to model and predict influenza A-H1N1 and B. The annual incidence of H3N2 and H1N1 between the 2002/2003 and 2017/2018 seasons in the United States (Figure 1, bottom panel) exhibit however a significant negative correlation (Spearman correlation of -0.64, p-value < 0.01), and H3N2 and B, a positive one (Spearman correlation of 0.58, p-value < 0.05). Additionally, we did not observe any novel antigenic change for either H1N1 or B in the last three years based on the PAV analysis (PAV < 0.1) (*4,6*). As a result, with a forecast of low H3N2 (median incidence rate 0.01) for the upcoming 2018/2019 season in the United States, we would expect a higher than average level for H1N1, and a lower than average level for B. For reference, the mean incidence rate of H1N1 was 0.10, and the mean value for B was 0.02 (based on the only two pandemic H1N1-dominant seasons of 2013/2014 and 2015/2016).

Although we correctly predicted an H3N2-dominant season for the past season that just ended, its observed incidence rate was more severe than our mean prediction (*2,5*). Also, the continuous model consistently predicts higher incidence rates than the cluster model, and has a tendency to overpredict (*1*). This indicates that there is room for improvement of EvoEpiFlu, to better incorporate information on antigenic change and to include other important information on vaccination and climate covariates. Future work is also needed to implement similar process-based dynamical models for H1N1 and for type B. Type B influenza in particular has been less studied, including the dynamics of its two lineages and their interaction with type A. Although the PAV values are low (<0.1) for both the Yamagata and Victoria lineages of influenza B, the CDC influenza surveillance system has captured novel antigenic variants in the Victoria lineage (*5,7*). Based on the observation that the Victoria lineage as a whole is declining (from 30% in the 2016-2017 season to 11% in the 2017-2018 season) among influenza B (*5,7*), we do not expect however an expansion and a high incidence rate for either the Victoria lineage or influenza B as a whole. Ideally, a multi-subtype model could be developed and evaluated for comprehensive forecasts of seasonal influenza in the United States, and potentially for better control and prevention.

## Data Availability Statement

All the data used in this study are from public databases and are publicly available to other researchers.

## Author Contributions

XD and MP designed the experiments; XD, YP, ML performed the experiments and analysis; XD and MP interpreted the results and wrote the manuscript.

## Acknowledgements

We acknowledge the National Supercomputer Centre at Guangzhou and the Research Computing Center at the University of Chicago for providing computational resources for this research. The authors are also very grateful to Tien Ming Lee, Yao-Qing Chen, Guanzheng Luo, Pei Xu and Huiying Zhao from Sun Yat-sen University, for inspiring discussions and suggestions, including during the earlier times of the formulation of the original model at UC.

## References

1. Du X, King AA, Woods RJ, Pascual M, Evolution-informed forecasting of seasonal influenza A (H3N2). Scinece Translational Medicine 2017, 9(413):eaan5325.

2. Du X, Pascual M, Incidence Prediction for the 2017-2018 influenza season in the United States with an evolution-informed model. Plos Currents Outbreaks 2018, January 17.

3. Shaman J, Karspeck A, Yang W, Tamerius J, Lipsitch M, Real-time influenza forecasts during the 2012-2013 season. Nature Communications 2013, 4:2837.

4. Du X, Dong L, Lan Y, Peng Y, Wu A, Zhang Y, Huang W, Wang D, Wang M, Guo Y, Shu Y, Jiang T, Mapping of H3N2 influenza antigenic evolution in China reveals a strategy for vaccine strain recommendation. Nature communications 2012, 3:709.

5. Gartem R, Blanton L, Elal AIA, Alabi N, Barnes J, Biggerstaff M, Brammer L, Budd AP, Burns E, Cummings CN, Davis T, Garg S, Gubareva L, Jang Y, Kniss K, Kramer N, Lindstrom S, Mustaquim D, O’Halloran A, Sessions W, Taylor C, Xu X, Dugan VG, Fry AM, Wentworth DE, Katz J, Jernigan D, Update: influenza activity in the United States during the 2017-2018 season and composition of the 2018-2019 influenza vaccine. MMWR Morb Mortal Wkly Rep 2018, 67(22):634–642.

6. Liu M, Zhao X, Hua S, Du X, Peng Y, Li X, Lan Y, Wang D, Wu A, Shu Y, Jiang T, Antigenic patterns and evolution of the human influenza A (H1N1) virus. Scientific Reports 2015, 5:14171.

7. Blanton L, Alabi N, Mustaquim D, Taylor C, Kniss K, Kramer N, Budd A, Gary S, Cummings CN, Chung J, Flannery B, Fry AM, Sessions W, Garten R, Xu X, Elal AIA, Gubareva L, Barness J, Dugan V, Wentworth DE, Burns E, Katz, J, Jernigan D, Brammer L, Update: influenza activity in the United States during the 2016-2017 season and composition of the 2017-2018 influenza vaccine. MMWR Morb Mortal Wkly Rep 2017, 66(25):668–676.

